# Nanotechnology-Driven Synergy: Research Effects of Curcumin Nanosuspension and Fluconazole Combination in Overcoming Azole Resistance in Candida Albicans

**DOI:** 10.1101/2025.04.25.650561

**Authors:** Cheng zhang, Jin Liu, Shoufu Sun

**Author notes:** Correspondence: Shoufu Sun. Cheng Zhang and Jin Liu have contributed equally to this work.

## Abstract

The increasing prevalence of invasive fungal infections, particularly life-threatening disseminated candidiasis caused by C. *albicans*, highlights the critical demand for novel therapeutic strategies. This study explores the antifungal potential of curcumin nanosuspension (CNS) alone and in combination with azoles, emphasizing synergistic mechanisms to counter drug resistance. CNS substantially improved curcumin’s aqueous solubility and enabled sustained in vitro release, overcoming its inherent bioavailability limitations. Checkerboard assays demonstrated synergistic interactions between CNS and fluconazole (FLC) against planktonic cells (FICI: 0.38–0.5) and early-stage C.*albicans* biofilms (FICI <0.5). Morphological analyses revealed that the combination suppressed hyphal formation and disrupted biofilm architecture—key virulence traits of C.*albicans*. Mechanistically, qPCR data indicated downregulation of adhesion-related genes (ALS1, ALS3, HWP1, EFG1), impairing fungal colonization. The synergy likely stems from dual mechanisms: facilitated FLC cellular uptake and inhibition of efflux pumps.These findings point out a nanotechnology-driven strategy to revitalize existing azoles through rational combination therapies. The improved bioavailability of CNS and their biofilm-penetrating capacity position this approach as a promising solution for managing refractory candidiasis, offering a pathway to circumvent antifungal resistance.

## INTRODUCTION

Candida *albicans* (C. *albicans*), a commensal fungal species commonly found in human microbiota, has emerged as one of the most clinically significant opportunistic pathogens in immunocompromised populations[1]. Clinical evidence indicates that this dimorphic fungus is responsible for both superficial and systemic infections, with particular clinical relevance in organ transplant recipients, AIDS patients, and chemotherapy patients where infection-associated mortality rates approach 40%[2-4]. Notably, the oral cavity serves as a primary reservoir for C. *albicans* colonization, with epidemiological studies demonstrating its isolation in 20%-50% of oral infection cases[5]. In immunocompromised hosts, this colonization frequently progresses to pathological manifestations including oral mucositis and candidiasis. Emerging research highlights the potential oncogenic implications of chronic C. *albicans* infections. Mothibe[6] reported significantly elevated fungal burdens in denture-wearing cancer patients compared to healthy controls, while Berkovits documented markedly increased C. *albicans* levels in oral squamous cell carcinoma patients relative to non-cancer populations[7]. These findings suggest a potential association between persistent fungal colonization and oral carcinogenesis that warrants further investigation. Of particular clinical concern is the propensity for localized oral infections to disseminate systemically. Given the escalating prevalence of immunocompromised populations worldwide and the emerging evidence of fungal oncogenic potential, the development of effective prevention and treatment strategies for oral candidiasis has become an urgent global public health priority.

The management of C. *albicans* infections relies heavily on effective antifungal agents. Current therapies primarily involve four major drug classes: azoles, amphotericin B, echinocandins, and fluconazole (FLC), with FLC being the most widely prescribed due to its cost-effectiveness and bioavailability[8, 9]. However, the emergence of azole-resistant strains, particularly in FLC-refractory cases linked to prolonged prophylactic use, complicates treatment[10]. This challenge is further intensified by the limited antifungal arsenal and the growing healthcare burden caused by C. *albicans*, underscoring the urgent need for novel therapeutic agents. In this context, natural products derived from Traditional Chinese Medicines (TCMs) have emerged as promising alternatives, offering advantages in bioavailability and reduced toxicity. Phytochemical studies have identified several TCM-derived compounds that enhance antifungal activity when combined with conventional azoles. Examples include berberine from Coptic chinensis and baicalein from Scutellarin baicalins, both of which exhibit significant effects through multi-target mechanisms[11, 12]. These findings highlight the potential of phytomedicine in addressing antimicrobial resistance.

Curcumin (CUR), isolated from turmeric, is renowned for its broad biological activities and non-toxicity, including antibacterial, antiviral, antifungal, and anti-inflammatory properties[13, 14]. Recent studies have further highlighted CUR’s antibacterial and antifungal potential against multidrug-resistant pathogens[15-17]. Combining natural compounds with conventional antimicrobials represents a promising strategy to combat fungal drug resistance[18]. For instance, Garcia-Goes demonstrated CUR’s ability to sensitize clinically resistant C. *albicans* strains to azole antifungals in vitro[19]. Despite these advantages, CUR’s clinical utility is hindered by physicochemical limitations such as poor water solubility, low bioavailability, rapid degradation, and inability to cross the blood-brain barrier[20]. Addressing these challenges particularly enhancing aqueous solubility and dissolution kinetics has emerged as a critical priority for translational applications.

Reducing the particle size of CUR is critical to enhancing its pharmaceutical performance[21]. Nanosuspensions, defined as submicron colloidal dispersions stabilized by minimal surfactant concentrations, offer significant advantages in drug delivery. When drug particles reach submicron dimensions (<1 μm), dissolution kinetics improve markedly due to increased surface area and enhanced saturation solubility[22]. These highly dispersed nano-systems enhance drug dissolution, absorption, and bioavailability across multiple administration routes[2, 23].Current pharmaceutical technologies allow the preparation of uniform and stable curcumin nanosuspensions(CNS). These optimized formulations address CUR’s solubility limitations by achieving a >40-fold increase in aqueous solubility and complete in vitro dissolution within 60 minutes[24]. Additionally, the modified-release characteristics of these nanosuspensions prolong therapeutic effects [25]. This technological advancement offers a strong experimental basis for overcoming CUR’s application barriers in antifungal therapy.

Studies have demonstrated that C. *albicans* exerts significant pathogenic effects on human hosts through its biofilm-forming capability and structured microbial communities’ adherent to both biotic and abiotic surfaces[26]. These biofilms substantially enhance fungal resistance to conventional antifungal agents and impair host immune defenses, thereby complicating clinical management of candidiasis and attracting considerable medical attention. The molecular mechanisms underlying biofilm formation involve coordinated expression of adhesion-related genes. Notably, the agglutinin-like sequence (ALS) gene family encodes cell-surface glycoproteins critical for host surface adherence[27]. Specifically, ALS1 encoded proteins mediate endothelial cell adhesion and participate in biofilm maturation[28]. Nails further identified ALS3 as a crucial regulator during initial hyphal development and biofilm establishment[29]. Complementary studies by Nobile revealed that HWP1 facilitates fungal-host cell adhesion while promoting biofilm structural integrity[30]. Additionally, Told demonstrated EFG1’s pivotal role in hyphal morphogenesis, phenotypic switching, and virulence regulation[31]. Furthermore, azole resistance mechanisms in C. *albicans* frequently involve overexpression of drug efflux pumps, representing a key pharmacological challenge in antifungal therapy [32].

This study focuses on addressing refractory and drug-resistant C. *albicans* infections. Utilizing a C. *albicans* biofilm model, we aim to prepare CNS and investigate the combined effects of CNS and FLC on biofilm formation. Further, we will employ qRT-PCR to assess the expression of adhesion-related genes, thereby elucidating the molecular mechanisms underlying the inhibition of C. *albicans* biofilms by the combined therapy. This research seeks to provide a feasible Nanotechnology therapeutic strategy for combating C. *albicans* infections, particularly those associated with biofilm-mediated resistance.

## MATERIALS AND METHODS

Three clinical C. *albicans* strains (CCA1, CCA2, and CCA3) were obtained from the Fungal Laboratory of Huashan Hospital, Fudan University (Shanghai, China). Among these, CCA3 was confirmed as a drug-resistant clinical strain by the Fungal Laboratory of Fudan University. Fungal cells were stored in Sabouraud dextrose agar (SDA) medium at -4°C and reactivated by subculturing on fresh SDA medium at 28°C for 48 hours prior to use. CNS were generously provided by the School of Pharmacy, Fudan University. Fluconazole (FLC) was purchased from Sigma Co., Ltd. Stock solutions of CNS (10.24 mg/mL) and FLC (2560 μg/mL) were prepared in distilled water and stored at -20°C until further use.

### *C. albicans* culture preparation and observation

Three clinical isolates of C. *albicans* (CCA1, CCA2, and CCA3) were employed in this study. Overnight cultures were prepared in Yeast Extract-Peptone-Dextrose (YPD) broth containing 1% (w/v) yeast extract, 2% (w/v) peptone, and 2% (w/v) glucose. Single colonies were inoculated into 30mL fresh YPD medium and incubated at 32°C with orbital shaking (160 rpm) for 15 hr. Cell pellets were obtained by centrifugation at 3,000 rpm for 10 min at 4°C, followed by two washes with sterile phosphate-buffered saline (PBS, pH 7.4). The washed cells were resuspended in RPMI-1640 medium (supplemented with L-glutamine, buffered with sodium bicarbonate) (Gibco, Grand Island, NY, USA) and adjusted to a final density of 2.0×10^6^ cells/ml using an improved Neubauer hemocytometer.

C. *albicans* cell suspensions at a concentration of 2×10^6^ CFU/mL were inoculated into sterile standard 96-well plates at 100μL per well and cultured for 2h, 6h, 12h, 24h, 36h and 48h, respectively. Following each culture period, 100μL of formaldehyde fixative solution was added to fix the cells for 20 minutes. Then, 0.001% crystal violet solution was added for staining, which lasted for 10 minutes. After staining, the plates were rinsed with phosphate-buffered saline (PBS) until the rinse solution remained clear. The biofilm morphology at the different time points was observed under an inverted phase-contrast microscope.Laser confocal microscopy was used to observe the viability of cells, scanning electron microscopy was employed to examine the morphological characteristics of C. *albicans* biofilms, and the XTT reduction assay was utilized to assess the metabolic activity of C. *albicans* biofilms.

### Preparation of CNS and analysis of its physical characterization

The procedure for preparing 200nm CNS was as follows: Accurately weigh 0.012g of curcumin and transfer it into an EP tube. Add 0.4mL of acetone and sonicate until the curcumin is fully dissolved to obtain the organic phase. Subsequently, stir the organic phase using a magnetic stirrer and transfer it to 4.6mL of 0.1% HPMC E5 solution. Remove the organic solvent with a rotary evaporator. Finally, adjust the 0.1% HPMC E5 solution to achieve a final curcumin concentration of 2.6 mg/mL in the preparation.

The prepared CNS was diluted 40-fold with ultrapure water. Then, 500μL of the diluted suspension was added to the sample tank, and the particle size and PDI were measured using a Malvern Zetasizer laser particle size analyzer at 25°C with a 60 s equilibrium time; each sample was measured three times, and the results were analyzed using Zetasizer Version 7.11. The zeta potential was measured using a zeta potential sample pool, with the diluted preparation added to ensure the liquid level exceeded the electrode level. For SEM observation, 50μL of the 40-fold diluted CNS was placed on a 50nm filter membrane, which was then dried, washed three times, naturally dried, cut to size, mounted on a sample table with conductive adhesive, and gold-sputtered for 60 s before imaging at 30,000× magnification and 5.0 kV emission voltage. To assess stability, 3 mL of CNS was kept at room temperature, and 200 μL samples were taken at 1, 2, 4, 6, 8, 10, 12, and 24 h, diluted 40-fold with ultrapure water, and analyzed for particle size and PDI using the laser particle size analyzer.

### XTT reduction assays

In the XTT reduction assays, menaphone solution (0.4 mM) was prepared by dissolving menaphone powder in acetone and stored at 4°C. XTT powder was mixed with PBS solution (pH 7.4) to obtain a 1 mg/mL test solution, which was stored at -70°C. Then, 2 μL of 0.4 mM menaphone solution, 40 μL of 1 mg/mL XTT solution, and 158 μL of PBS solution containing 200 mM glucose were thoroughly mixed and filtered through a 0.22 μm sterile filter. Biofilms of C. *albicans* strains, including the standard strain ATCC90028 and clinical strains CCA1, CCA2, and CCA3, were cultured for varying durations (2 h to 60 h). After removing the supernatant and rinsing twice with PBS, 200 μL of the XTT mixed solution was added to each well. The biofilms were then incubated in the dark at 37°C for 3 h. A microplate analyzer was used to detect the absorbance at 492 nm, which indicated the metabolic activity of the biofilms. All experiments were performed in triplicate and repeated three times.

### Scanning electron microscope (SEM) observation

In scanning electronic microscope (SEM) observation, C. *albicans* cell suspensions (2 × 10^6^ CFU/mL) were inoculated in RPMI-1640 medium in a 24-well plate with a cover glass at the bottom and incubated at 37°C for 48 h. Then, the cover glass was gently washed three times with sterile PBS buffer and fixed with 2.5% glutaraldehyde at 4°C for 72 h. After that, the fixed biofilm was washed three times with 0.1 mol/L phosphoric acid rinse solution (15 minutes each). Next, the biofilm was fixed with 1% osmium acid fixative for 3 h, followed by three more washes with the phosphoric acid rinse solution. The biofilms were dehydrated through a graded ethanol series (50%, 70%, 80%, 90%, and 100%, 20 minutes each). Then, the biofilm was immersed in a 1:1 mixture of 100% ethanol and ethyl acetate for 30 minutes, followed by soaking in ethyl amyl acetate for another 30 minutes. Finally, the dried biofilms were vacuum coated with gold and observed under a scanning electron microscope.

### Determination of minimum inhibitory concentrations(MIC)in planktonic cells

To determine the minimum inhibitory concentrations (MIC) in planktonic cells, we followed the micro-liquid-based dilution method per CLSI-M27A3 guidelines for antifungal susceptibility testing of CNS and FLC alone and in combination. For checkerboard assays, CNS and FLC concentrations ranged from 1–512 μg/mL and 1–32 μg/mL. C. *albicans* cell suspensions (2 ×10^3^ CFU/mL) were inoculated in RPMI-1640 medium in a 96-well plate, with different drug concentrations added to each well, including a blank control. After incubation at 37°C for 24 h in the dark, MIC_80_ was defined as the lowest drug concentration causing 80% growth inhibition compared to the blank group. MIC_80_ values were determined by measuring optical density at 492 nm (OD492) using a spectrophotometer (BioTek Synergy 2, USA). All experiments were performed in triplicate and repeated three times.

The interaction of FLC with CNS against C. *albicans* was evaluated using the fractional inhibitory concentration index (FICI). FICI was calculated as FICI = MIC (A in combination)/MIC (A alone) + MIC (B in combination)/MIC (B alone). Results indicated synergy when FICI ≤ 0.5, no interaction when 0.5 < FICI ≤ 4.0, and antagonism when FICI > 4.0.

### Determination of sessile minimum inhibitory concentrations(SMIC50) against C. *albicans* biofilms

The concentrations of CNS and FLC were set at 1–1024 μg/mL and 1–512 μg/mL. C. *albicans* cell suspensions (2×10^6^ CFU/mL) were inoculated into a 96-well plate containing RPMI-1640 medium. Different drug concentrations were added to each well, with a blank control group included. The plate was incubated at 37°C for 48 h in the dark. Biofilm formation was assessed using the XTT assay, and the OD492 was measured with a spectrophotometer (BioTek Synergy 2, USA). The sessile minimum inhibitory concentration (SMIC_50_) was defined as the lowest drug concentration causing 50% growth inhibition compared to the blank group. All experiments were performed in triplicate and repeated three times. The interaction of CNS and FLC on C. *albicans* biofilm was evaluated using the aforementioned FICI model.

### The morphology of biofilms was observed by Confocal laser scanning microscopy(CLSM) and scanning electron microscopy(SEM)

Confocal laser scanning microscopy (CLSM) was used to evaluate the inhibitory effects of drugs on C. *albicans* biofilms. The LIVE/DEAD Fungal Light Yeast Viability Kit (Molecular Probes, Eugene, OR, USA), containing Propidium Iodide (PI) and SYTO-9, was used to stain dead and live cells, respectively. C. *albicans* cell suspensions (2×10^6^ CFU/mL) were inoculated into laser confocal cell culture dishes containing RPMI-1640 medium. The final drug concentrations were as follows: 64 μg/mL CNS and 32 μg/mL FLC. The groups were set as: 1. Blank group; 2. FLC 32 μg/mL; 3. CNS 64 μg/mL; 4. FLC + CNS. After incubation at 37°C for 48 h, the PI, SYTO-9, and physiological saline were mixed in a ratio of 1.5 μL:1.5 μL:1000 μL, and 400 μL of the dye mixture was added to each dish. The dishes were then incubated at room temperature for 20 min in the dark. The maximum excitation/emission wavelengths for observation were 490/635 nm for PI and 480/500 nm for SYTO-9.

For further analysis, C. *albicans* cell suspensions (2×10^6^ CFU/mL) were inoculated into RPMI-1640 medium in 96-well and 24-well plates (with cover glasses). The same drug concentrations and grouping were applied. After incubation at 37°C for 48 h, biofilms in the 96-well plates were fixed with 100 μL of formaldehyde for 20 min, stained with 0.001% crystal violet for 10 min, and rinsed with PBS until clear. Biofilm morphology was observed under an inverted phase-contrast microscope. Biofilms in the 24-well plates were processed for SEM observation: the cover glasses were rinsed with sterile PBS, fixed with 2.5% glutaraldehyde, and then with 1% osmium acid fixative, dehydrated through an ethanol gradient, and critical-point dried. The dried biofilms were vacuum coated with gold and viewed under a scanning electron microscope.

### Quantitative real-time PCR (qRT-PCR) assay

The expression levels of biofilm-related genes (ALS1, ALS3, HWP1, and EFG1) were measured at 48 h post-drug treatment in ATCC90028 (1. Control group; 2. FLC 32 μg/mL; 3. CNS 64 μg/mL; 4. FLC + CNS) and CCA3 (1. Control group; 2. FLC 128 μg/mL; 3. CNS 256 μg/mL; 4. FLC + CNS). Cell walls were disrupted via liquid nitrogen grinding. Total RNA was extracted using 1 mL of Trizol Reagent. Then, 1 μL of RNA was used to synthesize cDNA. Real-time PCR was performed using a SYBR GREEN Mix, with gene-specific primers listed in Table 1. The relative RNA expression levels were calculated using the 2−ΔΔCt method.

**Table 1.**
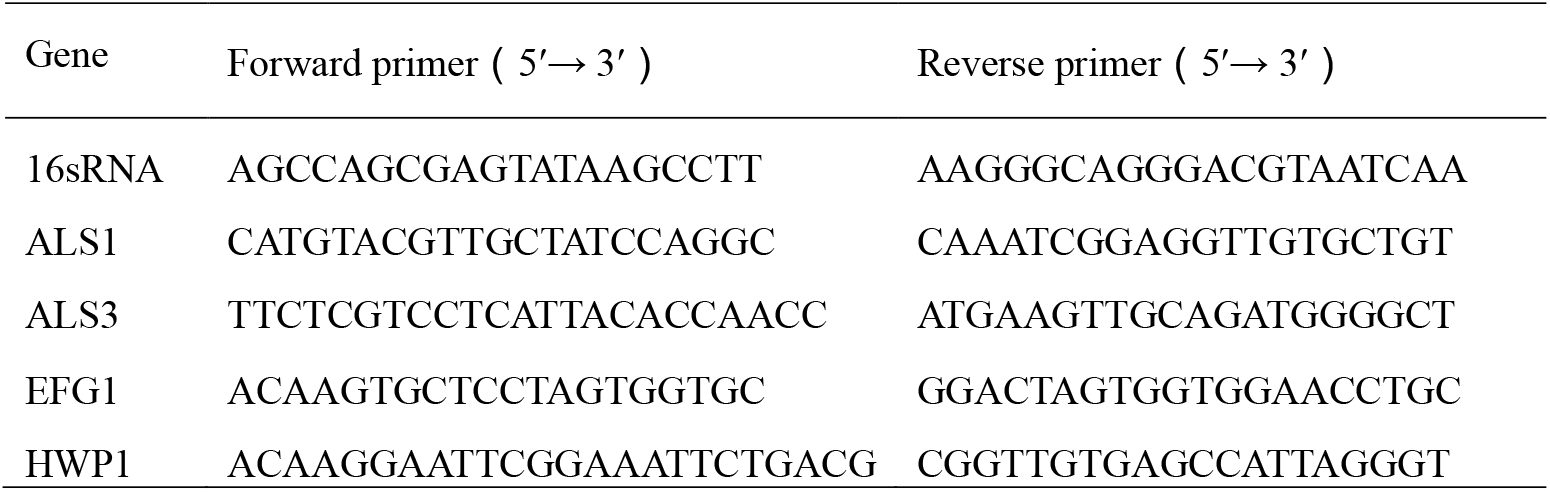
Real-time PCR was performed using a SYBR GREEN Mix, with gene-specific primers listed.

### Rhodamine 6G uptake and efflux assays

C. *albicans* strain CCA3 (FLC-resistant, 1 × 10^5^ CFU/mL) was cultured in YPD liquid medium at 35°C with agitation (180 rpm) for 18-19 h. Following incubation, cells were harvested by centrifugation (3,000 × g, 5 min) and washed three times with glucose-free phosphate-buffered saline (PBS). The cell density was adjusted to 1 × 10^7^ CFU/mL using a hemocytometer, followed by 2h de-energization in glucose-free PBS at 35°C with gentle shaking (50 rpm).

#### Uptake assay

Rh6G (10 μM final concentration) and CNS (128 μg/mL final concentration) were added to the cell suspensions. Control groups received Rh6G alone without CNS treatment. Intracellular fluorescence intensity was monitored at 10-min intervals using flow cytometry (BD FACSVerse™, excitation 488 nm, emission 530 nm) until signal stabilization occurred.

#### Efflux assay

De-energized cells were pre-incubated with 10 μM Rh6G at 35°C for 50 min, followed by immediate quenching in an ice-water bath (10 min) to arrest transport processes. After three cold PBS washes to remove extracellular dye, cells were resuspended in PBS containing either 128 μg/mL CNS or Rh6G alone (control). Fluorescence measurements were performed at 30-min intervals under flow cytometry until plateau values were obtained.

All experiments were conducted in triplicate using independent biological replicates. Data are presented as mean ± SD. Statistical significance was determined by two-tailed Student’s t-test (p < 0.05 considered significant).

### Statistical analysis

Statistical analyses were conducted using SPSS software (version 19.0; IBM, USA), with the significance level set at 5%.Normally distributed continuous variables were reported as mean ± standard deviation, and independent sample t-tests were used to compare the two groups. A value of p < 0.05 was considered statistically significant.

## RESULTS

### Formation of C. *albicans* biofilm

The formation of C. *albicans* biofilm was observed under an inverted phase-contrast microscope. Over time, the morphology evolved from yeast cells (2 h), to increased hyphae and pseudohyphae (6-12 h), followed by further elongation and proliferation of hyphae (24 h), and finally formed a dense network structure composed of the matrix, yeast cells, and numerous hyphae (48 h) (Figure 1A). The metabolic activity of the biofilms, as determined by the XTT reduction assay, gradually increased over time, peaking at 48 h (Figure 1B). SEM observation revealed a closely interwoven network structure of the biofilm, with true hyphal morphologies and pseudohyphae clearly visible (Figure 2A, B). Additionally, the poles of blastoconidial cells were identified as the primary budding sites, and variations in scar locations were also evident (Figure 2C, D).

**Figure 1.**
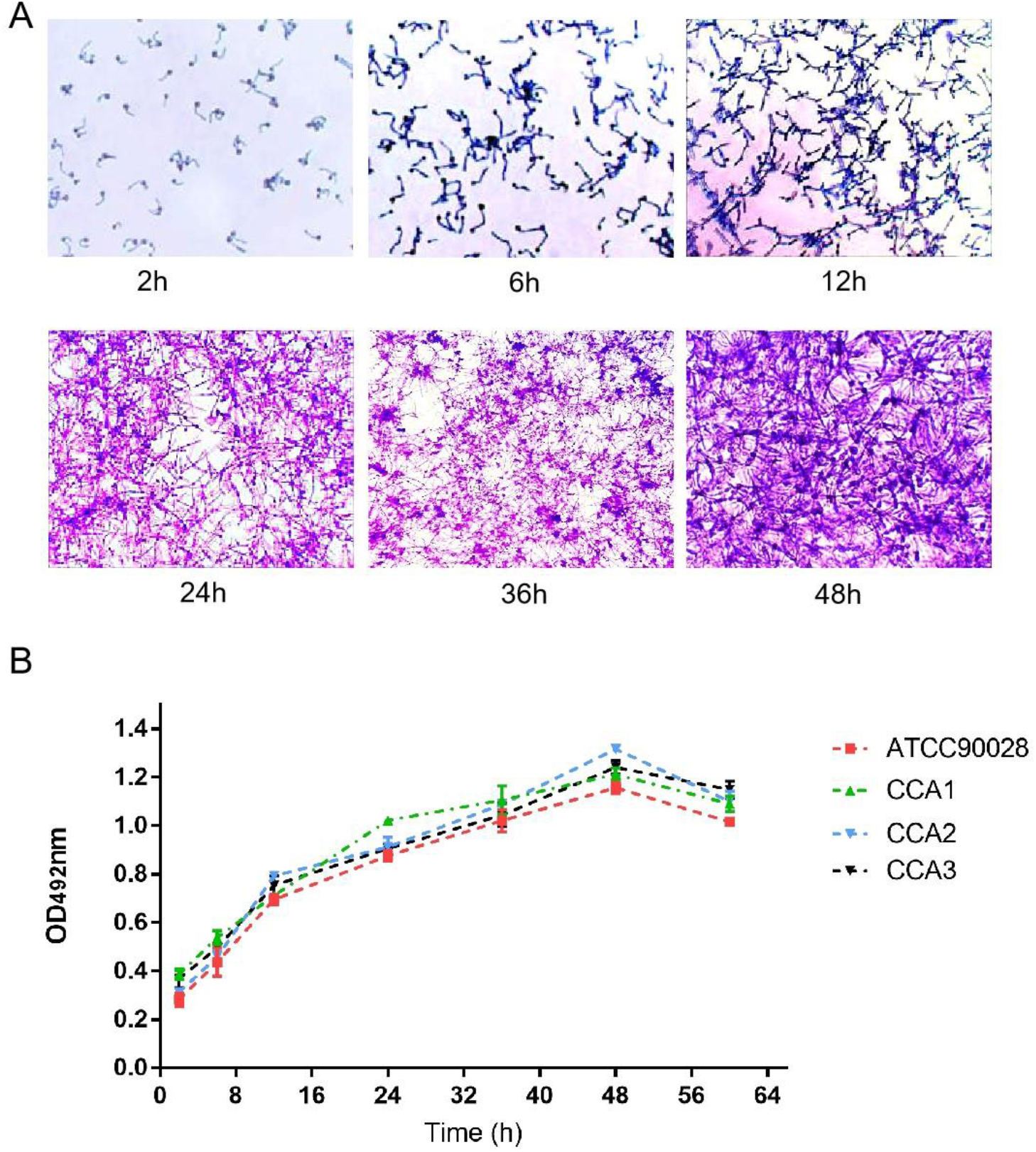
Formation and metabolic activity of C. *albicans* biofilms. (A) Biofilm formation was visualized under an inverted phase-contrast microscope. Crystal violet staining revealed biofilm development at various time points (2, 6, 12, 24, 36, and 48 h) in RPMI 1640 medium. (B) Metabolic activity of C. *albicans* biofilms was assessed using the XTT reduction assay. Data are presented as the mean ± standard deviation from two independent experiments conducted in triplicate.

**Figure 2.**
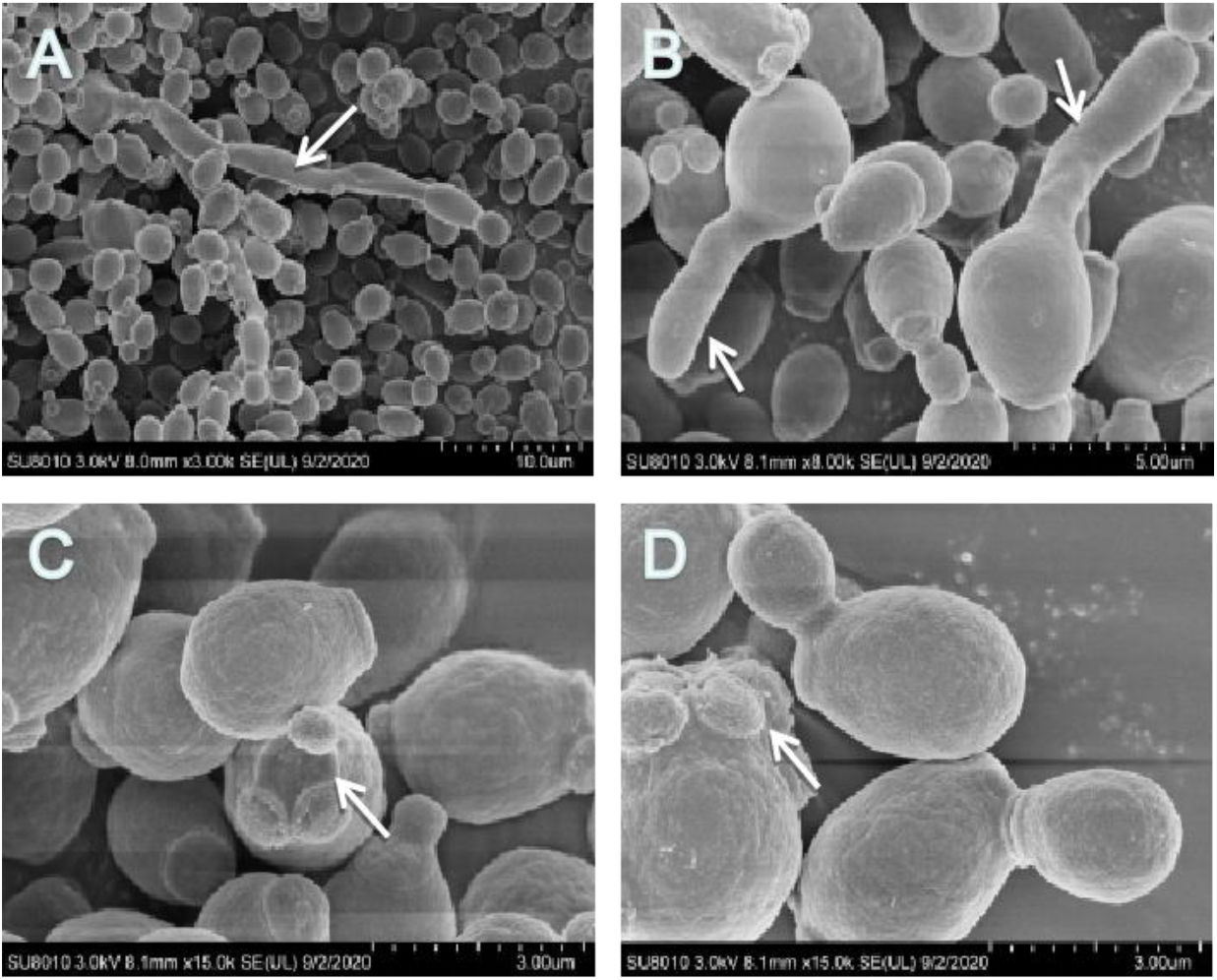
Scanning electron microscopy was used to observe the microstructure. (A) Biofilm structure: Mycelia form a network, arrange compactly, and cluster together. Note the true hyphal morphologies (open arrows). (B) Note the pseudohyphae structure. (C, D) Note the ring of scars (arrows) and a bud at one pole (tip) of the blastoconidial cell.

### Properties of Curcumin Nanosuspension

The particle size distribution of CNS, obtained via the described preparation process, is shown in Fig. 3A and B, indicating good dispersibility. The CNS had a particle size of 147.0 ± 1.304 nm, a PDI of 0.079 ± 0.013, and a zeta potential of -28.8 ± 0.251 mV. Electron microscopy (Fig. 3C) revealed that native curcumin particles were rod-shaped, micrometer-sized, and uneven in size, whereas CNS were spherical, smooth-surfaced, uniformly distributed, and non-aggregated. The particle size of the CNS, as observed via electron microscopy, was consistent with that measured by the laser particle size analyzer.

**Figure 3.**
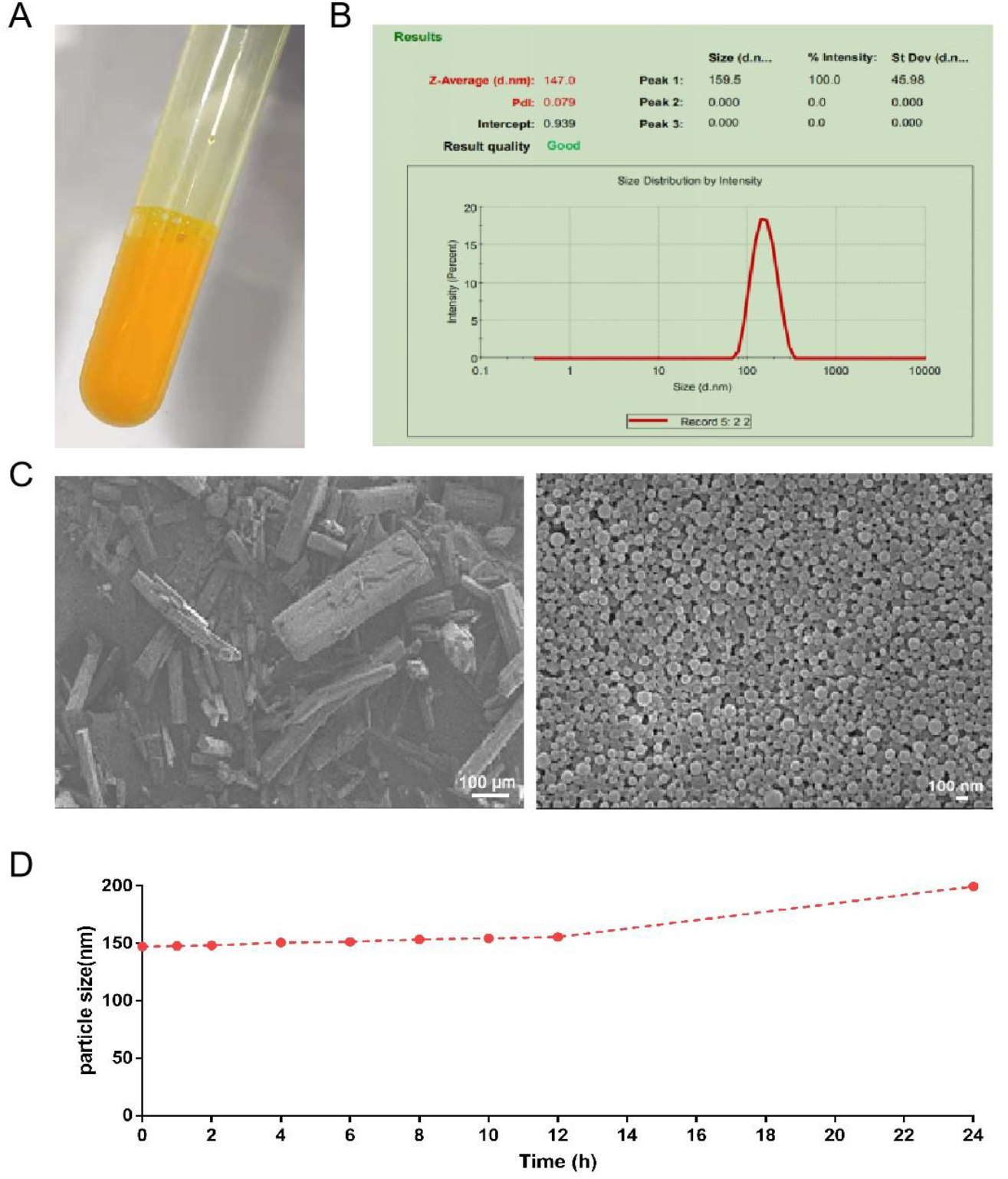
Preparation and Characterization of Curcumin Nanosuspension.(A) A mature and stable curcumin nanosuspension was prepared as described. (B) Particle size and PDI analysis via Malvern Zetasizer laser particle size analyzer revealed that CNS exhibited a particle size of 147.0 ± 1.304 nm, a PDI of 0.079 ± 0.013, and a zeta potential of -28.8 ± 0.251 mV. (C) Native curcumin particles were predominantly rod-shaped, micrometer-sized, and heterogeneous in size, whereas CNS were spherical, featured smooth surfaces, and showed uniform distribution. (D) Stability studies indicated that the particle size of the curcumin nanosuspension remained essentially unchanged over 24 hours.

The stability of the CNS was also evaluated (Fig. 3D). The particle size of CNS remained relatively constant for the first 12 hours and gradually increased to 200 nm after 12 hours of storage. These results suggest that the CNS prepared in this study exhibited good stability, providing a solid foundation for subsequent drug experiments.

### MIC and SMIC_50_ of C. *albicans*

Table 2 shows that CNS alone has no significant antifungal effect on C. *albicans* (MIC > 256 μg/mL). However, when CNS and FLC are used in combination against C. *albicans* planktonic cells, their MIC values are significantly reduced, with FICI values ranging from 0.25 to 0.5. These results indicate a strong synergistic effect between CNS and FLC against planktonic cells.

**Table 2.** MIC of CNS and FLC against Planktonic cells.

| Strains | Biofilm cells (μg/mL) |  |  |  |  |
| --- | --- | --- | --- | --- | --- |
|  | SMIC<sub>50</sub><br>Alone |  | SMIC<sub>50</sub><br>In combinationg |  | FICI<br>(Interaction) |
|  | CNS | FLC | CNS | FLC |  |
| ATCC90028 | 512 | 256 | 64 | 32 | 0.25 |
| CCA1 | 1024 | 512 | 128 | 64 | 0.25 |
| CCA2 | 1024 | 512 | 128 | 128 | 0.38 |
| CCA3 | 1024 | 1024 | 256 | 128 | 0.38 |

Further studies on the combined effect of CNS and FLC against C. *albicans* biofilms in four strains are presented in Table 3. CNS reduces the SMIC_50_ of FLC from 1024μg/mL to 32–128μ g/mL, with FICI values ranging from 0.25 to 0.38. This demonstrates a strong synergistic effect between CNS and FLC against all tested strains.

**Table 3.** SMIC_50_ of CNS and FLC against bioflim cells.

| Strains | Planktonic cells (μg/mL) |  |  |  |  |
| --- | --- | --- | --- | --- | --- |
|  | MIC<br>Alone |  | MIC<br>In combinationg |  | FICI<br>(Interaction) |
|  | CNS | FLC | CNS | FLC |  |
| ATCC90028 | 256 | 2 | 32 | 0.5 | 0.38 |
| CCA1 | 512 | 4 | 32 | 1 | 0.31 |
| CCA2 | 512 | 4 | 128 | 1 | 0.5 |
| CCA3 | 512 | 512 | 128 | 2 | 0.25 |

### Inhibition effects of CNS and FLC on C. *albicans* biofilms

SEM imaging of C. *albicans* ATCC90028 biofilms revealed distinct structural differences across treatment groups (Figure 4A). The blank control group displayed mature biofilms with interconnected hyphae forming a 3D network. Treatment with CNS or FLC alone reduced hyphae and bacterial counts, resulting in incomplete biofilm structures. However, the combination of CNS and FLC markedly decreased bacterial counts, with no visible hyphae or biofilm formation.

**Figure 4.**
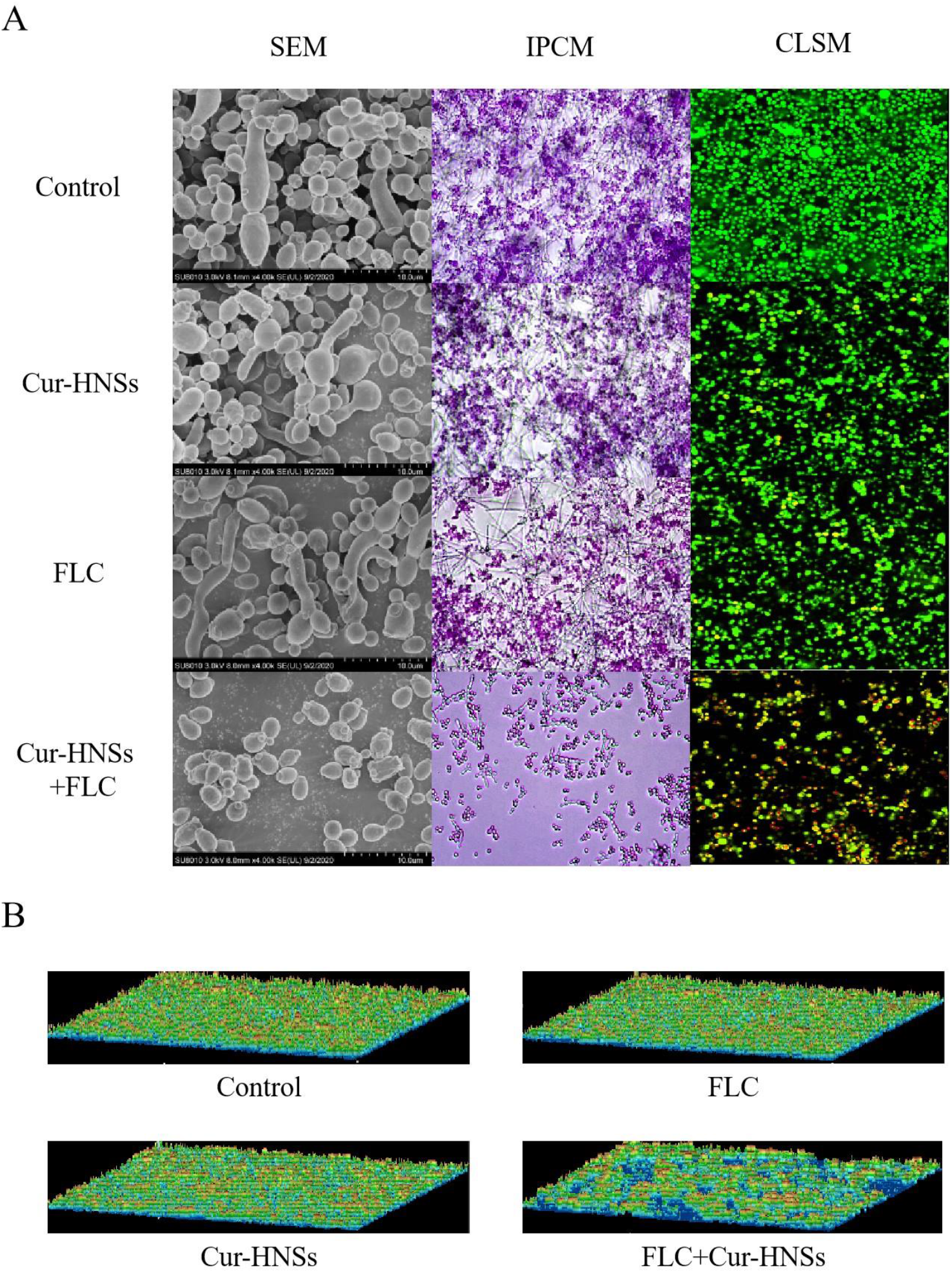
Effects of CNS and Fluconazole on C. *albicans* biofilm formation visualized by SEM and CLSM. (A) Control: adherent cells without drugs showing intact biofilm structures. (B) CNS (64 μg/mL): reduced biofilm formation. (C) Fluconazole (32 μg/mL): decreased biofilm density. (D) CNS + Fluconazole: significantly disrupted biofilm, few hyphae, and scattered cells.

Under an inverted phase-contrast microscope, the blank group showed dense hyphae forming a robust biofilm. Single-drug treatments altered the biofilm to a monolayer structure with reduced bacterial counts. In contrast, the combination group showed no biofilm structure, only isolated bacterial cells and minimal hyphae.

CLSM imaging further highlighted the synergistic effects of CNS and FLC. SYTO9 stained live cells green, while PI-stained dead cells red. The blank group displayed intense green fluorescence with no red fluorescence. Single-drug treatments reduced green fluorescence and introduced some red fluorescence, while the combination group showed sparse fluorescence with more red fluorescence, indicating enhanced inhibition.

3D reconstruction of biofilms using microscope software confirmed these observations. The blank and single-drug groups exhibited intact biofilm structures, whereas the combination group showed incomplete biofilm formation (Figure 4B).

### Quantification of adhesion and biofilm-related gene expression by qRT-PCR

The transcriptional levels of genes associated with adhesion and biofilm formation (ALS1, ALS3, HWP1, and EFG1) were quantified by qRT-PCR. All genes analyzed showed downregulation post - drug treatment. CNS or FLC alone caused slight downregulation compared to the control group. However, the combination of CNS and FLC led to significant downregulation, suggesting a notable impact on adhesion - related genes and hyphal growth (Figure 5).

**Figure 5.**
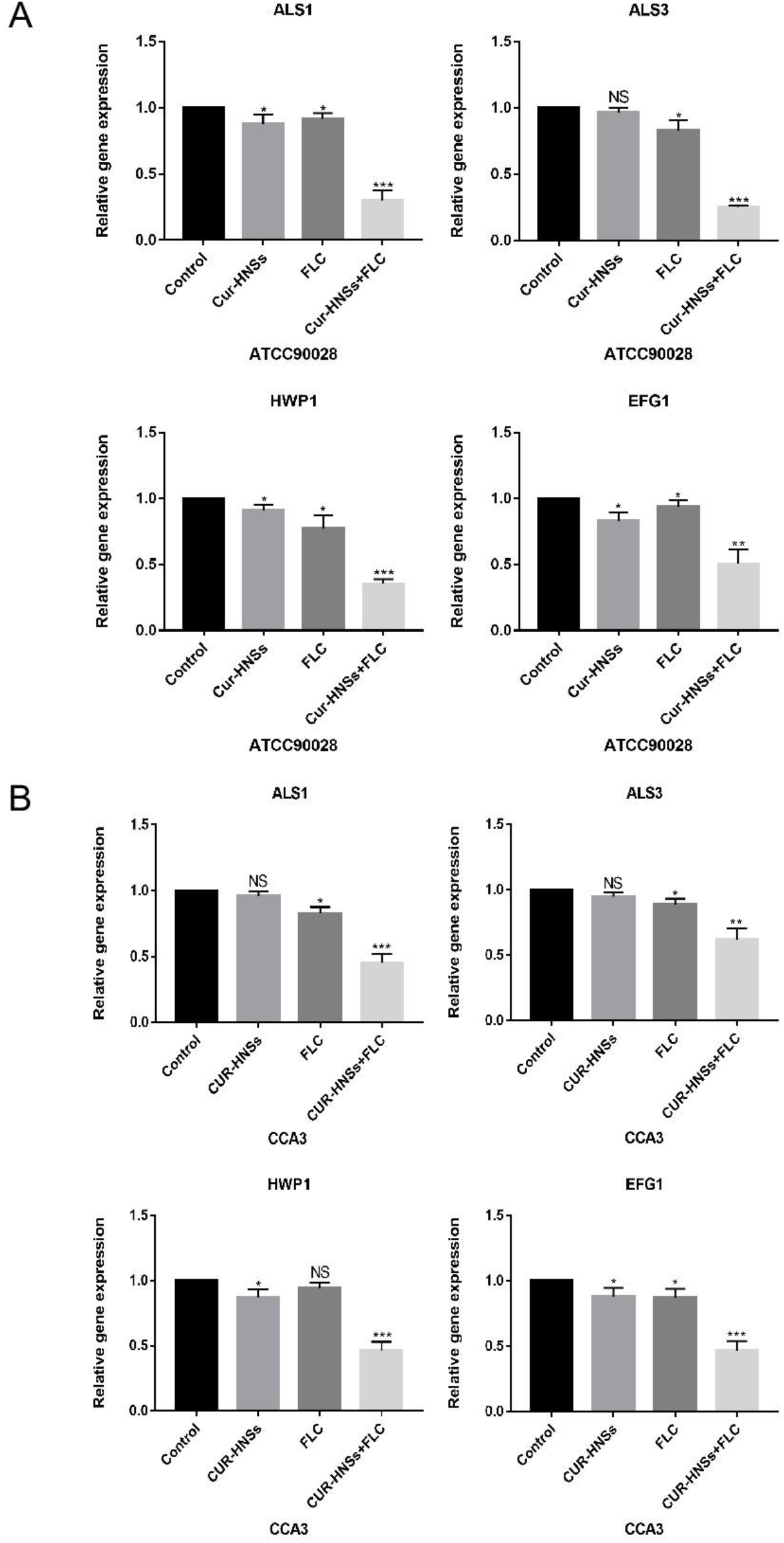
The relative expression of the genes (ALS1, ALS3, HWP1, EFG1) detected by real-time PCR. (A) ATCC90028: Control (biofilm-associated adherent cells without drugs), CNS (biofilm-associated adherent cells with CNS at the concentration of 64 μg/mL), Fluconazole (biofilm-associated adherent cells with fluconazole at the concentration of 32 μg/mL), Fluconazole + CNS (fluconazole at the concentration of 32 μg/mL+CNS at the concentration of 64 μg/mL). (B) CCA3: Control (biofilm-associated adherent cells without drugs), CNS (biofilm-associated adherent cells with CNS at the concentration of 256 μg/mL), Fluconazole (biofilm-associated adherent cells with fluconazole at the concentration of 128 μg/mL), Fluconazole+CNS (fluconazole at the concentration of 128 μg/mL + CNS at the concentration of 256 μg/mL). Results are mean ± standard error (M ± SE) from three independent experiments. *P* < 0.000, *P*< 0.01, *P* < 0.05, compared with the control cells.

### Uptake and Efflux of Rhodamine 6G

Both Rhodamine 6G (Rh6G) and FLC are substrates of drug transporter pumps located on the C. *albicans* cell membrane. In this study, Rh6G was used as a fluorescent alternative to FLC. Based on preliminary experiments showing that 128 µg/mL CNS significantly reduced the minimum inhibitory concentration (MIC) of FLC against FLC-resistant C. *albicans*, this concentration of CNS was selected for further investigation. As shown in Figure 6, a significant difference in intracellular fluorescence intensity was observed between the CNS group and the control group (P<0.05). This result indicated that the synergistic antifungal effects of FLC in combination with CNS against FLC-resistant C. *albicans* are related to the uptake and efflux mechanisms of FLC.

**Figure 6.**
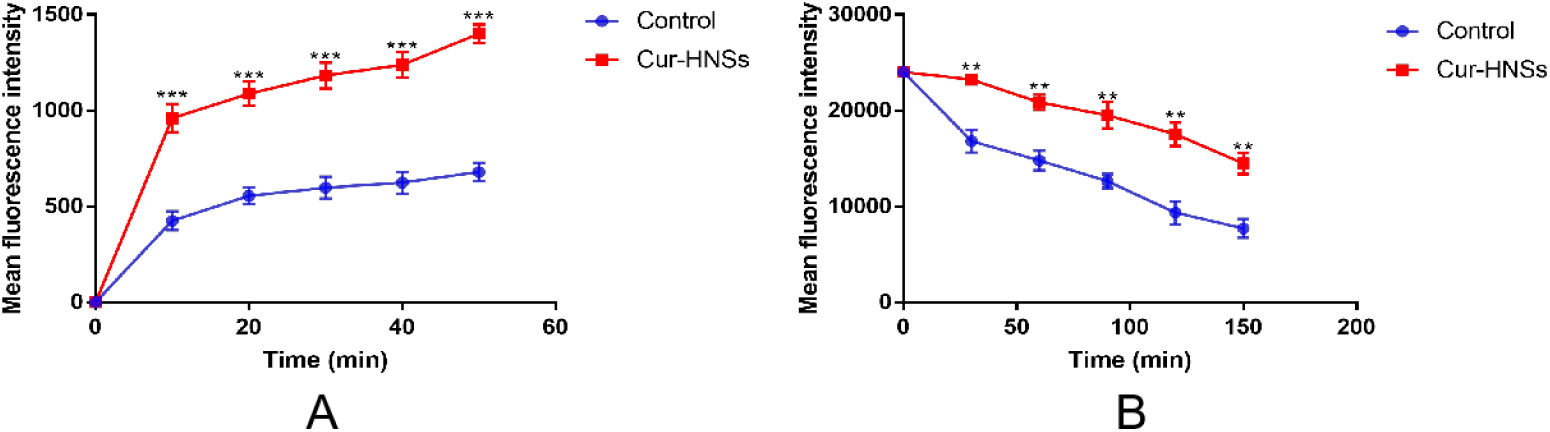
Effect of CNS on the uptake and efflux of Rh6G in *Candida albicans*. The uptake and efflux of Rh6G in the absence and presence of CNS (128 μg/mL) were determined by flow cytometry. Mean fluorescence intensity represents the intracellular Rh6G in *Candida albicans*. (A) The result shows the effect of CNS on Rh6G uptake. (B) The result shows the effect of GB on Rh6G efflux. Data were the means of three independent experiments.

## DISCUSSION

In recent decades, the incidence of fungal infections has risen rapidly, which can be largely attributed to the extensive use of broad - spectrum antibiotics, immunosuppressive agents, as well as medical implant devices. Among various fungal pathogens, C. *albicans* stands out as the most common one. It has the potential to induce a range of infections, from superficial epidermal ones to invasive and life - threatening conditions. This poses a particularly significant threat to immunocompromised patients’ FLC, an azole - class antifungal drug, is widely used for preventing and treating C. *albicans* infections thanks to its high efficacy and low toxicity. However, with prolonged and widespread use of FLC, a dramatic increase in C. *albicans*’ resistance to antifungal drugs, especially to FLC, has emerged as a major concern[33]. Furthermore, biofilms formed by C. *albicans*, which can adhere to both abiotic and biotic surfaces, serve as natural barriers against antifungal drugs and exhibit inherent resistance to most of them[34]. Considering these challenges, the development of novel treatment strategies to combat C. *albicans* resistance has become an urgent necessity.

The field of natural products derived from traditional Chinese medicinal herbs presents a rich reservoir for novel discoveries. Certain plant - based compounds exhibit potential as antifungal agents or synergistic partners with FLC. Curcumin demonstrates anti - C. *albicans* activity, yet its clinical application is hindered by poor solubility. Nanosuspension technology, through particle size reduction, has been utilized to improve its solubility, thereby enhancing its clinical potential. In recent years, combining traditional Chinese medicine monomers with antifungal drugs has proven to be an effective strategy for treating clinical drug - resistant strains. Combination therapy offers several benefits, including reduced drug dosages, minimized toxic and side effects, prevention of drug resistance development, and shortened treatment durations. In this study, it was discovered that CNS possess inhibitory effects against both C. *albicans* planktonic cells and biofilms. Furthermore, the inhibitory effects of the combination of CNS and FLC on C. *albicans* biofilms, as well as the underlying mechanisms, were investigated.

During the initial stage of biofilm formation (approximately 2 h), only round yeast - phase cells were observed under the microscope. These cells were relatively few in number and dispersed. At 12h of biofilm formation, mycelial structures became visible, and the cell population had increased. After 24 h of biofilm culture, a dense network of mycelial structures emerged, with cells aggregating into clusters and a substantial amount of extracellular matrix being secreted, thus preliminarily forming a biofilm - like structure. By 48 h, there was a further increase in mycelia, cells, and extracellular components. Following crystal violet staining, it was evident that a large number of mycelia had intertwined to form a network structure, characteristic of a typical biofilm. These findings aligned with those reported by other researchers[35].

The combined effects of the two drugs were evaluated using the checkerboard method. The fractional inhibitory concentration index (FICI) was employed to assess the interaction between them. The formula for FICI is as follows: FICI = MIC (A in combination)/MIC (A alone) + MIC (B in combination)/MIC (B alone). The smaller the FICI value, the stronger the synergy between the drugs. When FICI ≤ 0.5, the interaction is defined as synergy. The results indicated that CNS exhibited antibacterial activity against both standard and clinical strains of C. *albicans*. Moreover, when used in combination with FLC, CNS demonstrated significant synergistic antifungal activity, which considerably reduced the required dosage of FLC. Specifically, the SMIC50 of FLC was reduced from over 1024 μg/mL to 32 - 256 μg/mL. This synergistic anti - biofilm effect has the potential to overcome the resistance of C. *albicans* to FLC.

We observed the inhibitory effects of the combination of CNS and FLC on the formation of C. *albicans* biofilms using an inverted phase - contrast microscope, scanning electron microscope (SEM), and confocal laser scanning microscope (CLSM). Compared to the single - agent groups (CNS or FLC alone), the combination of the two drugs demonstrated a more pronounced impact on the biofilm structure. While CNS and FLC used individually could only thin the biofilm and reduce the formation of hyphal structures, the combined use of the two drugs led to a more significant disruption of the biofilm. Specifically, the combination destroyed the complete biofilm architecture, with only yeast - phase cells visible, indicating a more effective inhibition of biofilm development.

The Agglutin-like sequence (ALS) genes are crucial for the pathogenicity of C. *albicans*. Previous studies have shown that the ALS1 gene product mediates surface proteins that adhere to endothelial and epithelial cells during biofilm formation. The ALS3 gene encodes a cell surface protein that promotes hyphal formation and cell adhesion[36, 37]. The HWP1 gene encodes a protein that mediates the adhesion of C. *albicans* to epithelial cells and promotes biofilm formation[38]. Efg1 plays an extremely important role in the phenotypic transformation, hyphal formation, and virulence of C. *albicans*[39]. In our study, using RT-qPCR test, we observed that treatment with CNS or FLC alone could slightly downregulate the expression of four key biofilm - related genes (ALS1, ALS3, HWP1, and EFG1). However, when CNS were combined with FLC, the expression of these four genes was significantly reduced. This synergistic effect on gene expression inhibition likely contributed to the suppression of C. *albicans* biofilm formation.

The resistance of C. *albicans* to azole antifungal drugs, such as FLC, is often attributed to the increased activity of efflux pump proteins[40]. To explore whether the synergistic antifungal effects of FLC and CNS are related to the intracellular concentration of FLC, this experiment used Rhodamine 6G (Rh6G) as a fluorescent substitute for FLC. By observing the impact of CNS on intracellular Rh6G levels, we aimed to assess its effect on the drug transport pump activity of C. *albicans* strain CCA3.The results showed that CNS significantly promoted the intracellular absorption of Rh6G and reduced its efflux, indicating a marked inhibitory effect on the drug transport pump activity of CCA3. We speculate that CNS increases the intracellular concentration of FLC in CCA3 cells. This could reduce the MIC and SMIC50 values of FLC for CCA3, thereby reversing its resistance to FLC.

## CONCLUSION

This study demonstrates that CNS can effectively inhibit the growth of C. *albicans* and enhance the antifungal activity of FLC. Synergistic effects between the two drugs were observed through scanning electron microscopy and laser confocal microscopy. Mechanistic studies suggest that this synergy may result from the inhibition of hyphal growth, cell adhesion, and efflux pump activity. Further research is needed to fully elucidate the mechanisms underlying this synergistic effect.

## ACKNOWLEDGEMENTS

The authors would like to thank all the colleagues for their assistance in accomplishing this study.

